# Identification of a highly drought-resistant *pp7l hda6* mutant

**DOI:** 10.1101/2023.11.21.568111

**Authors:** Duorong Xu, Dario Leister, Tatjana Kleine

## Abstract

Plants have evolved efficient strategies to cope with drought stress, including stomata closure, significant changes in nuclear gene expression, and epigenetic mechanisms. Previously, we identified *Arabidopsis thaliana* PROTEIN PHOSPHATASE7-LIKE (PP7L) as an extrachloroplastic protein that promotes chloroplast development and high light, and salt tolerance. Here, we demonstrate that the *pp7l* mutant can withstand prolonged periods of drought stress. Interestingly, chloroplast development in *pp7l* recovers under drought conditions, despite growth of the mutant is impaired under normal growth conditions. To assess the (post)transcriptional changes occurring in the *pp7l* mutant under different durations of drought exposure, we used long non-coding RNA-sequencing. Compared to the previously reported drought-responsive changes in the wild type, the drought-responsive changes detected in the *pp7l* mutant were negligible. Our analysis of data generated in this study and previously motivated us to create a *pp7l hda6* mutant, which exhibits remarkable drought resistance. Notably, the growth penalty associated with *pp7l* was alleviated in the double mutant, ruling out a dwarf effect on the drought-tolerant trait of this genotype.

## 1 Introduction

Abiotic stress is one of the major constraints to global crop production and food security. Drought is caused not only by low precipitation, but also by temperature dynamics and light intensity, making it a key abiotic stress (Singh et al., 2018). Drought affects morphological, physiological, biochemical, and molecular characteristics of plants, with negative effects on photosynthetic capacity (Seleiman et al., 2021). Physiological changes may include the closure of stomata, changes in cell wall integrity, increased water use efficiency (WUE), and a reduction in relative water content (RWC), while biochemical changes include for example higher phytohormone production, in particular abscisic acid (ABA), and declined photosynthesis (Seleiman et al., 2021). However, a response to drought can have both positive and negative effects. Moreover, it is essential to consider that plant responses to drought involve a complex interplay of multiple factors, and understanding is far from complete. For example, stomatal closure prevents water loss by reducing transpiration, but also reduces the rate of photosynthesis by reducing CO_2_ uptake (Medrano et al., 2002; Lawson and Blatt, 2014).

On the molecular level, responses to drought and other stresses are accompanied by large-scale changes in nuclear (e.g., Bashir et al., 2019), and plastid (Xu et al., 2023) gene expression. In addition, in recent years, histone modifications have been shown to be among the epigenetic mechanisms that control gene expression in response to abiotic stress (Luo et al., 2017; Nunez-Vazquez et al., 2022), in particular drought (Li et al., 2021), by altering chromatin accessibility. One of the histone modifications is histone acetylation which involves the addition of acetyl groups to histone proteins mediated by histone deacetylases (HDACs). Acetylation is generally associated with relaxed chromatin structure, making the underlying genes more accessible for transcription. In *Arabidopsis*, HDA6 and HDA19 are the most extensively studied HDACs. Both participate in pathogen defense, regulation of flowering, senescence, and abiotic stress responses (Nunez-Vazquez et al., 2022), and both *hda6* (Kim et al., 2017) and *hda19* (Ueda et al., 2018) mutants show enhanced drought tolerance.

Here, we investigated the transcriptional behavior under drought of the *pp7l* mutant, which is devoid of PP7L, a member of a small subfamily of phosphoprotein phosphatases (Xu et al., 2019). We chose this mutant because, as we show here, it can survive very long periods of drought stress, and chloroplast development in *pp7l* recovers under drought, although growth of the mutant is impaired under normal growth conditions. We integrated our RNA-Seq analysis with previously published Col-0 RNA-Seq (Xu et al., 2023) and microarray (Kim et al., 2017) data, which motivated us to re-investigate drought tolerance of cell wall mutants, and to introduce a mutation in the histone deacetylase gene *HDA6* into *pp7l*. Strikingly, this led to the identification of an extremely drought-resistant *pp7l hda6* mutant in which the growth retardation seen in *pp7l* plants is relieved.

## 2 Materials and Methods

### 2.0 Plant material and growth conditions

All Arabidopsis (*Arabidopsis thaliana*) lines used in this study share the Columbia genetic background. Seeds were sown on 1/2 MS medium containing 1% (w/v) sucrose and 0.8% (w/v) agar, incubated at 4°C for 2 days, and transferred to a climate chamber under a 16-h-light/8-h-dark cycle with a light intensity of 100 μE m^-2^ s^-1^ at 22°C. For physiological experiments, the plants were grown on potting soil (A210, Stender, Schermbeck, Germany) for 3 or 4 weeks.

The mutants analyzed in this study included *pp7l-1, pp7l-2, cry1cry2, cop1-4*, which were used previously (Xu et al., 2019), *hda6-7* (N66154; also known as *rts1-1*) (Aufsatz et al., 2002) and *sal1* (SALK_020882) (Wilson et al., 2009). The T-DNA line SALK_201895 (*hda6-11*) was obtained from the Arabidopsis Biological Resource Center. The homozygous *hda6-11, pp7l-1 hda6-11* and *pp7l-2 hda6-11* mutant lines were identified using primers suggested by the SALK Institute (http://signal.salk.edu/cgi-bin/tdnaexpress). The position of the T-DNA insertion in *HDA6* was confirmed by sequencing and lies +4 bp from the start codon of the *AT5G63110* gene.

For drought treatments, 7-d-old seedlings were transferred to pots, and grown for a further 2 weeks under normal growth conditions. Three-week-old plants were subjected to drought conditions by withholding water for the indicated times. The drought phenotypes were documented with a camera (Canon 550D, Krefeld, Germany).

### 2.1 Stomatal aperture measurement

Mature stomata of epidermal strips from 3- to 4-week-old plants were used for stomatal aperture measurements. After dark adaptation for 24 h, the stomata of *pp7l* plants were found to be significantly open (see Supplementary Figure S1). Thus, all of the plants were initially kept in darkness for 48 h, and were then illuminated with 30 μmol m^-2^ s^-1^ blue light, 50 μmol m^-2^ s^-1^ red light or 50 μmol m^-2^ s^-1^ far-red light for 2 days. Epidermal strips of leaves were peeled off the abaxial side of the leaf under dim light using forceps. To examine drought-mediated stomatal closure, epidermal strips were peeled from plants growing under normal or drought conditions. Stomatal apertures were photographed under a microscope (Zeiss, Oberkochen, Germany) and measured with the help of ImageJ software.

### 2.2 Chlorophyll *a* fluorescence measurement

Chlorophyll fluorescence parameters were measured using an imaging Chl fluorometer (Imaging PAM, M-Series; Walz, Effeltrich, Germany). The method employed involved dark adaptation for 30-min, followed by the determination of Fv/Fm. F_0_ was measured at a low frequency of pulse-modulated measuring light (4 Hz, intensity 3, gain 3, damping 2), while Fm was quantified following saturation pulses of approximately 2700 μmol photons m^−2^ s^−1^ for 0.8 s. The calculations and plotting of the parameters were performed using the ImagingWinGigE software.

### 2.3 In vivo labeling of chloroplast proteins

Chloroplast protein labeling was performed essentially as described previously (Xu et al., 2019) with the following modifications. For in vivo labeling, 100 mg young leaves were incubated for 30 min in the labeling buffer containing 20 μg/mL cycloheximide, 10 mM Tris-HCl, 5 mM MgCl_2,_ 20 mM KCl, pH 6.8, and 0.1% (v/v) Tween 20, to block the synthesis of nuclear-encoded proteins. Then, [^35^S]Met (1 mCi) was added to the same solution. After labeling for 30 min under light conditions of 80 μmol photons m^−2^ s^−1^, the leaves were washed and frozen in liquid nitrogen. Subsequently, total proteins were isolated and subjected to SDS-PAGE. Radiolabeled proteins were visualized by autoradiography (Typhoon™ laser scanner, GE Amersham).

### 2.4 Nucleic acid extraction

Leaf tissue (100 mg) was homogenized in extraction buffer containing 200 mM Tris/HCl (pH 7.5), 25 mM NaCl, 25 mM EDTA and 0.5% (w/v) SDS. After centrifugation, DNA was precipitated from the supernatant by adding isopropyl alcohol. After washing with 70% (v/v) ethanol, the DNA was dissolved in distilled water.

### 2.5 RNA-sequencing (RNA-Seq)

Total RNA from plants was isolated using TRIzol Reagent™ (Thermo Fisher Scientific, Waltham, MA, USA) and purified using an RNA Clean & Concentrator (Zymo Research, Irvine, USA) according to the manufacturer’s instructions. RNA integrity and quality were assessed with an Agilent 2100 Bioanalyzer (Santa Clara, USA). Ribosomal RNA depletion, generation of RNA-Seq libraries and 150-bp paired-end sequencing on an Illumina HiSeq 2500 system (Illumina, San Diego, USA) were conducted as described (Xu et al., 2023). Three independent biological replicates were used per condition (0, 6, 9, 12 and 15 days). Sequencing data have been deposited to Gene Expression Omnibus (Edgar et al., 2002) under the accession number GSE202931.

### 2.6 Chloro-Seq and 3D RNA-Seq analysis

To detect editing and splicing efficiencies of chloroplast-encoded transcripts, a modified Chloro-Seq pipeline (Malbert et al., 2018) was used as described (Xu et al., 2023).

To quantify overall transcript accumulation, the 3D RNA-Seq pipeline (Guo et al., 2021) was used. To this end, RNA-Seq reads were prepared on the Galaxy platform (https://usegalaxy.org/). Adaptors were removed with Trimmomatic (Bolger et al., 2014), and sequencing quality was accessed with FastQC (http://www.bioinformatics.babraham.ac.uk/projects/fastqc/). Transcript abundances were calculated using Salmon (Patro et al., 2017) and AtRTD2-QUASI as reference transcriptome (Zhang et al., 2017). The generated files were uploaded into the 3D RNA-Seq app (https://3drnaseq.hutton.ac.uk/app_direct/3DRNAseq; (Calixto et al., 2018; Guo et al., 2021) and transcript per million reads (TPMs) were calculated using the implemented lengthScaledTPM method. Weakly expressed transcripts and genes were filtered out based on the mean-variance trend of the data. Relative expression values were calculated as described (Xu et al., 2023).

### 2.7 Determination of Gene Ontology (GO) enrichments

GO enrichments were obtained from the Database for Annotation, Visualization and Integrated Discovery (DAVID; (Huang da et al., 2009), applying a cut-off of 5-fold enrichment compared to the expected frequency in the Arabidopsis genome and an FDR (Benjamini-Hochberg) ≤ 0.05.

## 3 Results and discussion

### 3.0 The *pp7l* mutant can survive long periods of drought stress – also under strictly controlled water conditions

In *Arabidopsis thaliana* (hereafter, Arabidopsis), loss of PP7L, a member of a small subfamily of phosphoprotein phosphatases (PPPs), results in pleiotropic phenotypes including decreased resistance to salt, high-light and temperature stress (Xu et al., 2019; Xu et al., 2020b). During further phenotypical characterization, we noticed that the stomata of *pp7l* mutant seedlings were constitutively open under blue, red, and far-red light, as well as in darkness (Supplementary Figure 1A), and the width-to-length ratio of the stomatal aperture was significantly higher under all conditions (Supplementary Figure 1B). This is reminiscent of the stomatal behavior of the *cop1-4* (*constitutively photomorphogenic1-4*) mutant (Mao et al., 2005). Moreover, isolated *cop1-4* leaves treated with abscisic acid (ABA), as well as *cop1-4* soil-grown plants exposed to drought conditions, do not close their stomata, and accordingly, water loss from detached *cop1-4* leaves is higher than from wild-type leaves. Nevertheless, soil-grown *cop1-4* copes better with drought than the wild type (Moazzam-Jazi et al., 2018). To test whether *pp7l* mutants also displayed altered tolerance to drought, water was withheld for 15 days from 3-week-old Col-0 and *pp7l-1* mutant plants.

Interestingly, upon re-watering of these plants for three days, 60% of them survived, while all Col-0 plants died (Supplementary Figure 1C). In addition, when water was withheld for 24 days, again all Col-0 plants died, while all *pp7l-1* plants survived (Supplementary Figure 1D). However, *pp7l* is a dwarf mutant (Xu et al., 2019). Thus, it is possible that *pp7l* and other dwarf mutants are more resistant to drought conditions because their roots extract water from the soil at a lower rate. To investigate this in more depth, we performed a second drought experiment in which we exposed Col-0 and *pp7l-1* to the same drought conditions by controlled watering through adjusting pot weights (Figure 1A). Because chlorophyll fluorescence measurements are a noninvasive and rapid method to assess the impact of drought stress on the functionality of photosystem II (PSII) (Chen et al., 2016; Hu et al., 2023), phenotypes and maximum quantum yield of photosystem (PS) II (Fv/Fm) were determined. Remarkably, Col-0 plants died after 46 days of drought treatment, whereas the *pp7l-1* mutant still displayed photosynthetic activity as indicated by the measurement of Fv/Fm (Figure 1A). In another experimental set-up, we included the dwarf mutants *cop1* (higher water loss under drought; Moazzam-Jazi et al., 2018) and *sal1* (no higher water loss of detached rosettes under drought; Wilson et al., 2009) as controls. Because COP1 likely acts downstream of the cryptochrome (CRY) pathway as a repressor of stomatal opening, and the non-dwarf *cry1 cry2* mutant is also drought tolerant (Mao et al., 2005), this double mutant was also included in our study. Water was withheld from 3-week-old plants and pictures were taken after 16 and 24 days. All mutants appeared to be drought-tolerant after 16 days. They accumulated less anthocyanin and leaves looked healthier than those of the WT (Figure 1B). The remarkable ability of the *pp7l* mutant to cope with drought stress was again underpinned after 24 days. After this period, *pp7l* showed no signs of wilting, while *sal1* had begun to wilt, and *cop1-4* and *cry1 cry2* were clearly wilting. Analysis of epidermal stomata from the abaxial side of mature and young WT and *pp7l* leaves showed that *pp7l* stomata had a higher width-to-length ratio than WT under normal conditions (Figure 1C and D). Strikingly, mature leaves of *pp7l* were able to close their stomata just as well as mature and young leaves of the WT even under drought exposure. However, stomata of young *pp7l* leaves were indifferent to drought stress and showed the same open appearance as under control conditions (Figure 1C).

**Figure 1.**
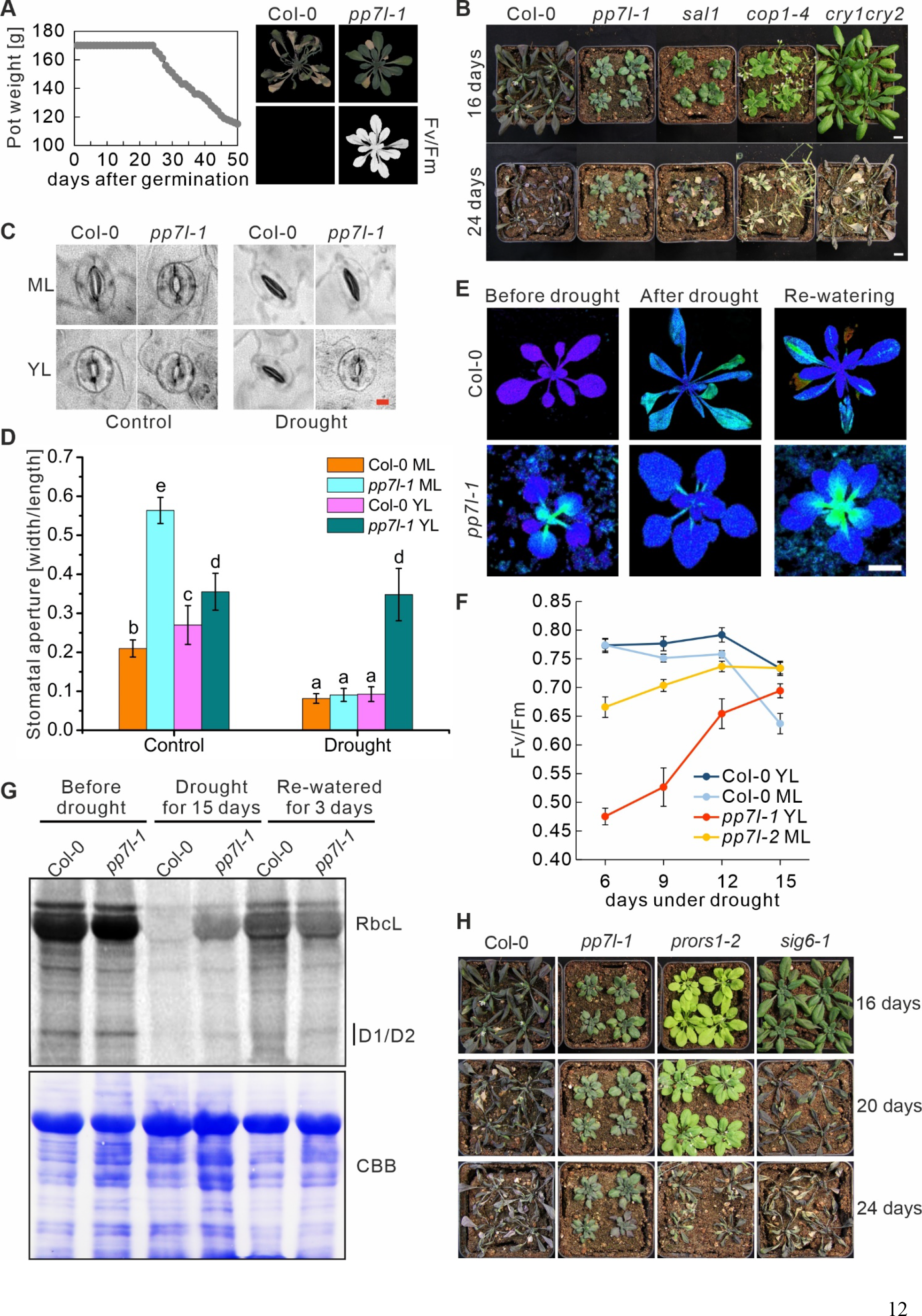
Soil-grown *pp7l* plants are more drought tolerant than the wild type. **(A)** The water status was controlled by watering pots to the same pot weight. After 24 days pot weight was gradually reduced. 46-day-old Col-0 plants died while *pp7l* plants were still viable as indicated by detectable photosynthetic activity (maximum quantum yield of photosystem II, Fv/Fm). **(B)** Col-0 and the different genotypes were grown for 3 weeks under control conditions, and then water was withheld for 16 and 24 days. Bar = 1 cm. **(C)** Representative images of stomata of Col-0 and the *pp7l-1* mutant under control conditions and after drought exposure for 12 days. Bar =10 μm. **(D)** Analysis of stomatal apertures of Col-0 and *pp7l-1* plants treated as in panel C. Each column represents the mean value of data for three biological replicates. Bars show standard deviations and lowercase letters (a to e) indicate independent classes according to Student’s t-test with a significance level of *P* < 0.001. **(E)** Recovery of chloroplast development in *pp7l* plants following drought stress documented by Imaging-PAM pictures of 3-week-old *pp7l* plants taken before and after exposure to drought stress induced by withholding water for 15 days, and after re-watering for 3 days. **(F)** Graph displaying F_v_/F_m_ values of 3-week–old Col-0 and *pp7l* mutant plants subjected for the indicated time to drought stress. The data represent mean values ± SDs of three independent experiments each containing 8 plants. YL, young leaves; ML, mature leaves. **(G)** In vivo pulse labeling of chloroplast proteins with [^35^S]Met in the presence of cycloheximide indicates that translation occurs at higher rates in *pp7l*-*1* chloroplasts under drought stress, compared to lower rates when plants were re-watered. Proteins were resolved by SDS-PAGE after pulse-labeling for 30 minutes, and visualized by autoradiography. The Coomassie-Brilliant-Blue (CBB)-stained membrane served as a loading control. **(H)** Col-0 and the different genotypes were grown for 3 weeks under control conditions, and then water was withheld for 16, 20 and 24 days.

In conclusion, the hypothesis of differential water usage by the dwarf mutants cannot be dismissed, however we can classify *pp7l* as a bona fide survivor of long drought periods.

### 3.1 Chloroplast development recovers in *pp7l* plants during drought stress

Chloroplast development in young leaves is delayed in plants lacking functional PP7L under normal growth conditions (Xu et al., 2019; Figure 1E), and the difference in stomatal behavior between young and mature *pp7l* leaves is noteworthy (see Figure 1C). To determine whether photosynthetic performance was also differentially regulated during drought, Fv/Fm was measured in young and mature leaves after 6, 9, 12 and 15 days of drought stress. Fv/Fm of WT leaves remained relatively constant (at around 0.77) for up to 12 days of drought exposure; after 15 days it had dropped to 0.74 and 0.63 in young and mature leaves, respectively (Figure 1F). Surprisingly, but in agreement with the healthier appearance of *pp7l*, the low Fv/Fm values measured in *pp7l* leaves (0.47 and 0.66 for young and mature leaves, respectively) rose during the drought-stress period to 0.69 and 0.74, respectively, showing a pronounced recovery (especially of young leaves) after 15 days of drought exposure (Figure 1E and F). During the re-watering period, Fv/Fm values of *pp7l* leaves resembled the low values observed for *pp7l* plants grown under continuous, well-watered conditions. Previously it was shown that chloroplast translation is reduced in young *pp7l* mutant seedlings (Xu et al., 2019). Therefore, we investigated the effect of drought on the translation of chloroplast transcripts by in vivo labeling, which enables the quantification of de novo protein synthesis. To this end, synthesis of cp-encoded proteins was monitored by pulse labeling of WT and *pp7l*-*1* mutant plants in the presence of cycloheximide, which inhibits the translation of nuclear-encoded proteins. This was done before and during drought stress, as well as after re-watering for 3 days. After pulse labeling for 30 minutes, the de novo synthesis of RbcL, D1, and D2 proteins was observed to be increased in *pp7l*-*1* compared to WT under drought conditions. However, the synthesis of these proteins was reduced in *pp7l-1* compared to WT after 3 days of re-watering (Figure 1G). To test whether mutants in cp gene expression are generally more drought tolerant, we tested the *prors1* and *sig6* mutants. These mutants are impaired in organellar translation (Pesaresi et al., 2006) and transcription (Ishizaki et al., 2005), respectively, and were subjected to drought stress. The WT wilted after 16 days, while the *prors1* and *sig6* mutants wilted after 20 and 24 days, respectively (Figure 1H). However, the *pp7l* mutant remained healthy throughout the whole period.

We conclude that – in contrast to the WT – chloroplast development in the *pp7l* mutant is positively influenced by drought stress. Additionally, a disturbance in chloroplast gene expression does not generally lead to a drought tolerance as observed when PP7L is lacking.

### 3.2 Drought stress and its impact on the *pp7l* chloroplast (post)transcriptome

Prompted by the long survival and recovery of Fv/Fm in the *pp7l* mutant under drought stress which might be interesting for both organellar and nuclear RNA expression patterns, RNA sequencing was performed in a way that allowed to detect nuclear as well as organellar transcripts. However, it should be noted that tRNAs cannot be reliably identified because their small size and many modifications make them difficult to amplify using either the RNA-Seq method used in this publication or conventional mRNA-Seq protocols. RNA was isolated from 3-week-old pp7l plants grown under optimal conditions (time point 0 d; 0d_DS) and after water deprivation for 6, 9, 12, and 15 d (6d_DS, 9d_DS, 12d_DS, and 15d_DS, respectively) as described previously (Kim et al., 2017) and grown in parallel with previously characterized Col-0 (Xu et al., 2023). The experiment was repeated separately with different batches of plants, resulting in three biological replicates per time point, with a sequencing depth of approximately 25 million 150-bp paired-end reads for each of the samples (Supplementary Table 1).

Because of the strong effect of drought stress on the Fv/Fm phenotype of *pp7l* plants, we first investigated accumulation of chloroplast (cp) transcripts under drought stress. A gene was considered to be differentially expressed (DEG) if it showed an absolute log_2_ fold change ≥1 (≥ 2-fold linear change) in expression in at least one contrast group (adjusted *P* < 0.05; Supplementary Table 2). To illustrate expression changes, heatmaps of Z-means were generated for transcripts that showed expression changes in at least one condition examined (Figure 2A). First, expression changes between *pp7l* and Col-0 under normal growth conditions (time point 0) were analyzed. Of the 71 protein-coding cp genes that were above the detection threshold, five transcripts were elevated and only one reduced which was that of *psbK* encoding PSII reaction center protein K. Notably, in 4-day-old *pp7l* seedlings grown under well-watered conditions, cp transcripts also generally accumulated to higher levels than those observed in wild-type controls (Xu et al., 2019). In addition, three, four, 18, and 17 transcripts were at least 2-fold reduced (compared to the 0d_DS sample) in *pp7l* plants after 6, 9, 12 and 15 days of drought, respectively, while no transcript was induced. However, the relatively constant – and transiently even higher, although not 2-fold – expression of genes encoding the PSII reaction center proteins PsbA, PsbC, and PsbD in *pp7l* plants throughout drought stress is noteworthy (Supplementary Table 2). The most affected gene that was already down-regulated after 6 days codes for MatK, the only cp-localized splicing factor, and which is involved in splicing of several *tRNA*s, *atpF, rpl2* and *rps12-2* (de Longevialle et al., 2010). Genes whose transcripts were reduced during the later time points code for example for the F, G, and H subunits of the NAD(P)H dehydrogenase complex, the PSII subunits K and E, the PSI subunits J and C and ribosomal protein S15.

**Figure 2.**
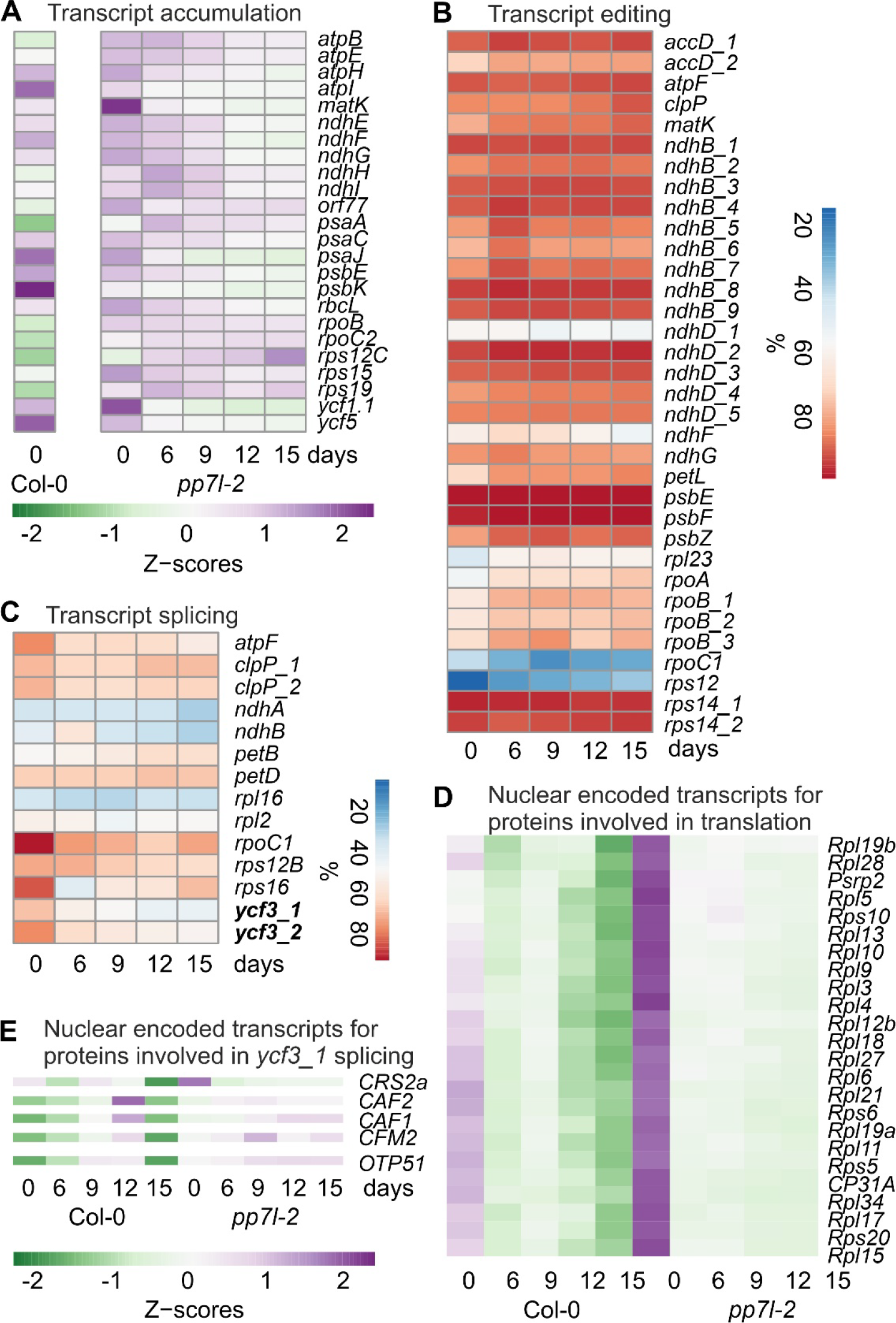
Impact of drought stress on accumulation, editing and splicing of chloroplast transcripts. Water was withheld for 15 days from 3-week-old *pp7l* plants grown under standard conditions (0 days), and RNA-Seq was performed as described in Materials and Methods on RNA extracted from plants harvested at the indicated time-points. **(A)** Heatmap illustrating accumulation of selected chloroplast transcripts (Z-scores) during the drought time-course. Low to high expression is represented by the green to purple transition. Note that Z-scores are calculated for each individual transcript over the time course. Col-0 data were retrieved from (Xu et al., 2023). **(B, C)** Percentages of editing **(B)** and splicing **(C)** events during the time-course. **(D, E)** Heatmap illustrating accumulation of nuclear encoded transcripts for proteins involved in chloroplast translation **(D)**, and *ycf3_1* splicing **(E)** during the drought time-course.

To examine post-transcriptional changes in cp transcripts, editing and splicing efficiencies were calculated. Editing efficiency was not notably diminished, and in some cases slightly enhanced (Figure 2B). However, splicing efficiency was reduced under drought stress for 5 out of 14 transcripts (Figure 2C). Interestingly, this is in contrast to Col-0, where splicing efficiency was not significantly decreased after 15 d of drought stress, but in some cases was slightly increased (Xu et al., 2023). With the exception of MatK, all splicing and editing factors are encoded in the nuclear genome, and chloroplast ribosomes are a mosaic of proteins encoded by both the cp and the nucleus. To investigate a putative correlation between nuclear gene expression of chloroplast splicing factors and the respective splicing events, transcript accumulation of genes encoding proteins involved in chloroplast gene expression were determined. Here, also transcript accumulation of Col-0 under drought was incorporated. Plotting their Z-scores and sorting the transcripts into clusters according to their expression performance under drought, resulted in two clusters (Supplementary Figure 2). One cluster showed a transient up-regulation of transcripts, and a final down-regulation after 15 d in Col-0, while these transcripts did not change or were slightly up-regulated in *pp7l*. In the other cluster transcript accumulation was higher in *pp7l* under well-watered conditions than in Col-0, but transcripts decreased under drought in both Col-0 and *pp7l* (Figure 2D, Supplementary Figure 2).

This cluster contains genes encoding mainly ribosomal proteins, which is intuitively in contrast to the improved cp translation in *pp7l* under drought stress (see Figure 1G). In addition, there appears to be no simple correlation between the accumulation of nuclear-encoded cp transcript maturation factors and cp post-transcriptomic changes. This is also due to complex interactions of splicing factors. For example, in land plants, intron 1 of *ycf3* (*ycf3_1*) is spliced by at least CRM Family Member 2 (CFM2) and RNA splicing factor CRS2 together with CRS2-associated factors CAF1 and 2 (de Longevialle et al., 2010), but only *CRS2* transcripts are decreased under drought (Figure 2E). Intron 2 of *ycf3* is spliced by ORGANELLAR TRANSCRIPT PROCESSING 51 (OTP51) (de Longevialle et al., 2010), whose transcripts are slightly enhanced under drought (Figure 2E), whereas splicing of *ycf3_2* is reduced (Figure 2C).

In conclusion, there is no simple correlation between mRNA levels of nuclear-encoded genes affecting cp gene expression and cp (post)-transcriptome changes under drought.

### 3.3 Behavior of cell wall mutants under drought conditions

We also used the generated RNA-Seq data to compare the overall nuclear gene expression behavior of *pp7l* under drought with that of Col-0. Hereafter, a gene was considered a DEG if it showed an absolute log_2_ fold change ≥1 in expression in at least one contrast group and a more stringent adjusted *P* of < 0.01. Slightly more genes were down-(54.4%) than up-regulated in *pp7l*, but the differences in gene expression increased only moderately when compared to the previously generated Col-0 dataset (Figure 3A; Supplementary Table 3). This is also reflected in a principal component analysis (PCA) of gene expression data across the samples, which showed that drought (64% of variance) was the main driver of gene expression in Col-0, but not in *pp7l* (Figure 3B).

**Figure 3.**
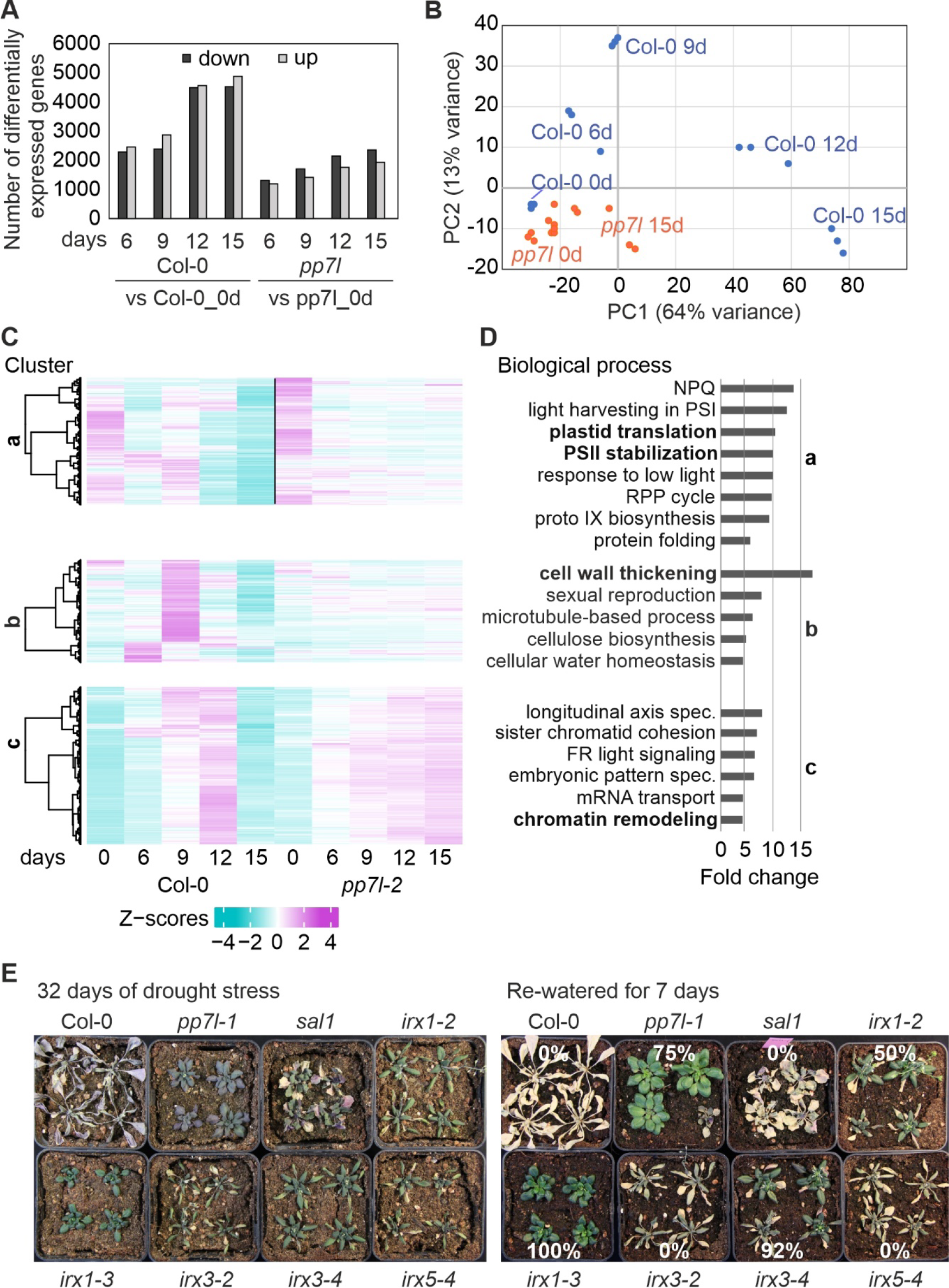
Overview and gene ontology analysis of gene expression changes under drought. **(A)** Numbers of differentially expressed genes (absolute log_2_ fold change ≥1; in at least one contrast group with an adjusted *P* < 0.01) in Col-0 and *pp7l* plants subjected to drought exposure compared to the respective control (0 days). **(B)** Principal Component Analysis (PCA) plot visualizing variation between genotypes and time-points based on RNA-seq data. **(C)** Hierarchical clustering was used to partition the differentially expressed (DE) genes into clusters with the Euclidean distance and ward.D clustering algorithm. **(D)** Graphs illustrating non-redundant Gene Ontology (GO) term enrichment for the biological process category according to DAVID (Huang da et al., 2009). GO terms with a ≥5-fold change and a Benjamini-corrected *P*-value of <0.05 are shown. LHC, light-harvesting complex; NPQ, nonphotochemical quenching; RPP, reductive pentose-phosphate; PS, photosystem. **(E)** Col-0 and the different genotypes were grown for 3 weeks under control conditions, water was withheld for 32 days, and then re-watered for 7 days.

The kinetic data were further explored by sorting the *pp7l*-related transcript changes into the nine different clusters which were previously identified in the Col-0 drought set-up (Xu et al., 2023), and three clusters caught our attention. Cluster a contains the genes that were strongly down-regulated in Col-0 at 12d_DS and 15d_DS, but these genes are only moderately repressed in *pp7l*. Gene Ontology (GO) analysis of this cluster showed that the enriched GOs in the biological process (BP) category included those related to chlorophyll biosynthesis, cp translation, and photosynthesis-related processes (Figure 3C and D). This underscores the greater susceptibility of Col-0 to drought stress.

In cluster b, genes assigned to the GO categories “cellular water homeostasis”, together with “cell wall thickening” (more than 15-fold enrichment) are specifically induced in Col-0 after 6 or 9 days of drought stress, and decline under progressive drought, but are only slightly changed in *pp7l*. It was previously found that disruption of *CELLULOSE SYNTHASE GENE 8* (*CESA8*; *IRREGULAR XYLEM 1, IRX1*) enhances drought and osmotic stress tolerance in Arabidopsis (Chen et al., 2005). Inspection of the cell wall thickening genes in cluster b showed that *IRX1* was indeed included, as well as *IRX3, IRX5* and *IRX9*. IRX1, 3 and 5 are involved in cellulose synthesis of the secondary cell wall (Taylor et al., 2003), and it was noted before that a defect in IRX1, IRX5 (Hernandez-Blanco et al., 2007) or IRX3 (Xu et al., 2020a), results in the up-regulation of ABA-responsive genes.

However, how individual CESA/IRX proteins are organized within the CSC remains unclear. It has been suggested that CESA7/IRX3 has a highly restricted position within the CSC, and in contrast, CESA8/IRX1 appears to have very low class specificity (Kumar et al., 2017). To observe the drought response of *irx1* mutants under our drought conditions, to confirm that other *irx1* mutant alleles are also drought tolerant, and to clarify whether disruption of *IRX3* or *IRX5* can confer drought tolerance or that of IRX3 can even lead to higher drought tolerance, we observed previously identified *irx1-2, irx1-3, irx3-2, irx3-4*, and *irx5-4* mutants (Xu et al., 2020a) in comparison to *pp7l* and *sal1* mutants when water was withheld for 32 days and re-watered for 7 days. After re-watering, WT, *sal1, irx3-2*, and *irx5-4* plants died (Figure 3E). However, *irx1-2, pp7l, irx3-4*, and *irx1-3* plants had survival rates of 50%, 75%, 92%, and 100%, respectively. Of note is that this variation among alleles was previously observed when studying the performance of cell wall mutants under conditions that restrict chloroplast translation (Xu et al., 2020a).

In summary, drought stress induces a massive transcriptional reprogramming of transcripts for cell wall protein transcripts, but of the cell wall proteins investigated, specifically IRX1 deficiency in *irx1-3* reliably confers drought tolerance.

### 3.4 The double mutant *pp7l hda6* is extremely drought resistant

Kim et al. (2017) identified acetate as a driver of drought resistance, based on the observation that, under prolonged drought stress, larger amounts of *PDC1* and *ALDH2B7* transcripts accumulated in the drought-tolerant *histone deacetylase 6* (*hda6*) mutant than in the control line DR5 (Kim et al., 2017; Figure 4A). Both of these genes code for enzymes in the acetate biosynthesis pathway. In our set-up, the degree of induction of *PDC1* and *ALDH2B7* was higher in Col-0 than in DR5. In drought-exposed *pp7l* plants, these transcripts were also induced, but at levels lower than that seen in Col-0 (Figure 4A). This contrasts sharply with the effect of the *hda6* mutation compared to the DR5 control. The third cluster (cluster c) that caught our attention contained genes that behaved similarly to *PDC1* and *ALDH2B7* in the *hda6* mutant. This cluster comprises genes whose transcript levels transiently increased in Col-0 and dropped after prolonged drought, but remained up-regulated in *pp7l* from 9 days of drought stress on. Strikingly, the GO term “chromatin remodeling” is enriched in this cluster, and *HDA6* itself shows such an expression profile, together with *HISTONE DEACETYLATION COMPLEX 1* (*HDC1*; Figure 4B and C). Note here the differential accumulation of *HDC1* isoforms (Figure 4C). HDC1 interacts with HDA6 and HDA19 to facilitate histone deacetylation (Perrella et al., 2013). Moreover, levels of both *HDA6* and *HDC1* mRNAs were already elevated in *pp7l* relative to Col-0 under normal growth conditions. This finding was unexpected, because theoretically, this would imply that HDA6 and its interactor are required to *withstand* drought. Hence, introduction of a *HDA6* mutation into *pp7l* would be expected to abrogate *pp7l*’s drought-tolerant phenotype, although the *hda6* single mutant is drought tolerant. To test this, we created *pp7l hda6* double mutants (Supplementary Figure 3), and withheld water from 3-week-old Col-0, *hda6, pp7l, pp7l hda6* and – as a control – *sal1* plants. After 18 days, Col-0 plants had begun to die, and their Fv/Fm values were very low (Figure 4D). In contrast, the younger leaves of *hda6, sal1* and *pp7l* plants were still able to photosynthesize and did not wilt, corroborating earlier reports of their drought resistance phenotypes. Moreover, the *pp7l hda6* double mutant outperformed all other mutants, displaying the highest photosynthetic capacity, low anthocyanin accumulation and no signs of wilting. Notably, the dwarf phenotype of *pp7l* was also suppressed in the *pp7l hda6* double mutant. Consequently, a dwarf effect on the drought-tolerant phenotype of this genotype under our experimental conditions can be excluded.

**Figure 4.**
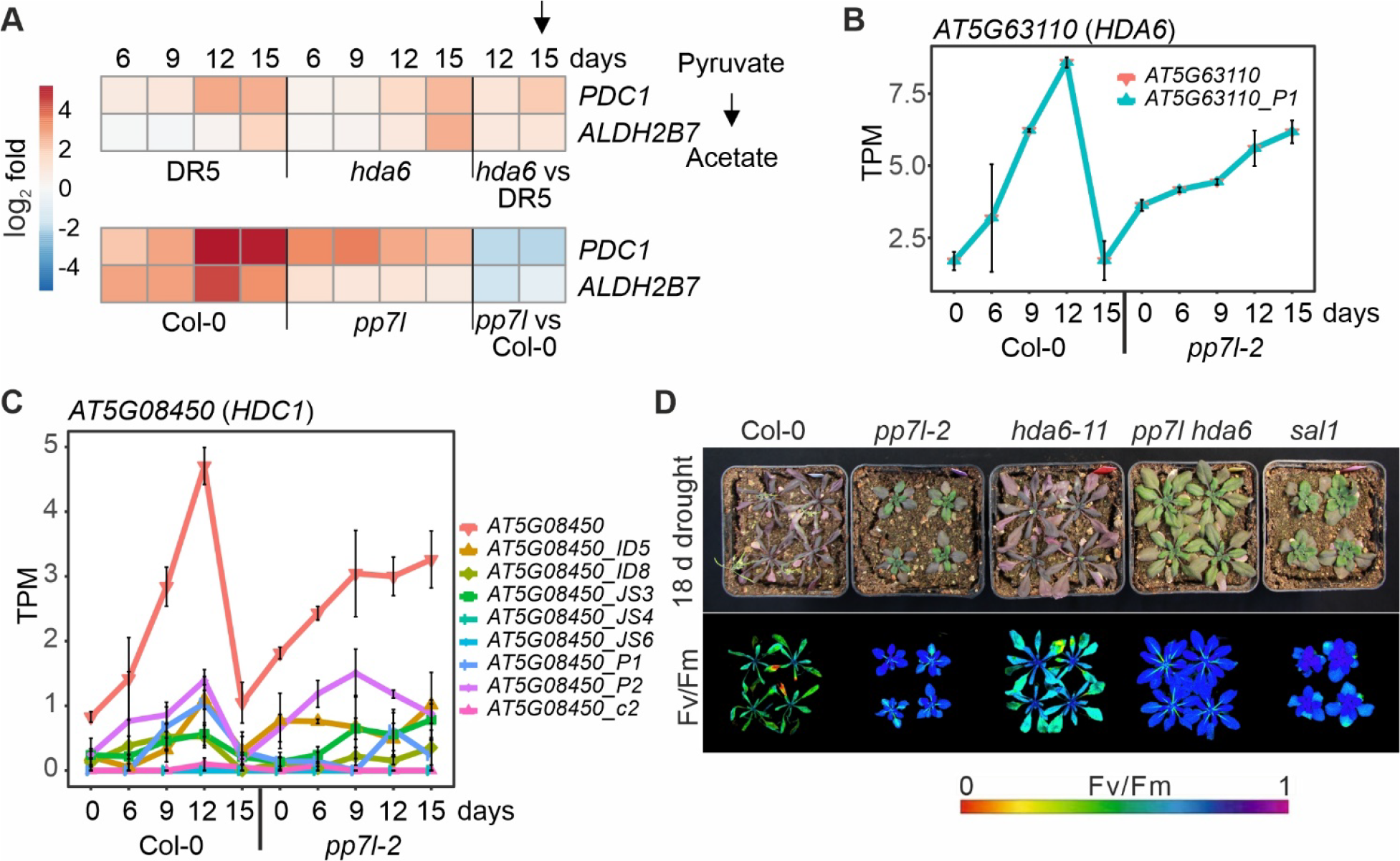
The *pp7l hda6* mutant is extremely drought resistant. **(A)** Heatmap illustrating the log_2_ fold changes compared to time-point 0 days of pyruvate decarboxylase *PDC1* and acetaldehyde dehydrogenase *ALDH2B7* transcripts in the experiment conducted by Kim et al. (2017) (upper panel) and in this study (lower panel). **(B, C)** Expression profiles of *HDA6* **(B)** and *HDC1* **(C)** at the whole-gene level and at the level of detected transcript isoforms. PS, percentage of expressed transcripts spliced; TPM, transcripts per million reads. **(D)** Col-0 and mutant plants were grown in separate pots in the same container, and water was withheld from 3-week-old plants for 18 days.

These results show that both PP7L and HDA6 constrain drought tolerance, and that loss of both proteins results in an extremely drought-resistant mutant in which growth penalty of *pp7l* has been released.

## 4 Conflict of Interest

The authors declare that the research was conducted in the absence of any commercial or financial relationships that could be construed as a potential conflict of interest.

## 5 Author Contributions

Conceptualization, T.K. and D.X.; Methodology, D.X., and T.K.; Formal Analysis, T.K.; Investigation, D.X., and T.K.; Resources, D.L. and T.K.; Writing – Original Draft, T.K.; Writing – Review & Editing, D.X., D.L., and T.K.; Funding Acquisition, D.L. and T.K.; Supervision, T.K.

## 6 Funding

This work was supported by the Deutsche Forschungsgemeinschaft (TRR175, Project C01 to T.K. and Project C05 to D.L.).

## 7 Acknowledgments

We thank Elisabeth Gerick for excellent technical assistance.

## 8 Data Availability Statement

Sequencing data have been deposited to Gene Expression Omnibus (Edgar *et al*., 2002) under the accession number GSE202931.

## Notes

### Competing Interest Statement

The authors have declared no competing interest.

## References

Aufsatz, W., Mette, M.F., van der Winden, J., Matzke, M., and Matzke, A.J. (2002). HDA6, a putative histone deacetylase needed to enhance DNA methylation induced by double-stranded RNA. EMBO J 21(24), 6832–6841. doi: 10.1093/emboj/cdf663.

Bashir, K., Matsui, A., Rasheed, S., and Seki, M. (2019). Recent advances in the characterization of plant transcriptomes in response to drought, salinity, heat, and cold stress. F1000Res 8. doi: 10.12688/f1000research.18424.1.

Bolger, A.M., Lohse, M., and Usadel, B. (2014). Trimmomatic: a flexible trimmer for Illumina sequence data. Bioinformatics 30(15), 2114–2120. doi: 10.1093/bioinformatics/btu170.

Calixto, C.P.G., Guo, W., James, A.B., Tzioutziou, N.A., Entizne, J.C., Panter, P.E., et al. (2018). Rapid and Dynamic Alternative Splicing Impacts the Arabidopsis Cold Response Transcriptome. Plant Cell 30(7), 1424–1444. doi: 10.1105/tpc.18.00177.

Chen, Y.E., Liu, W.J., Su, Y.Q., Cui, J.M., Zhang, Z.W., Yuan, M., et al. (2016). Different response of photosystem II to short and long-term drought stress in Arabidopsis thaliana. Physiol Plant 158(2), 225–235. doi: 10.1111/ppl.12438.

Chen, Z., Hong, X., Zhang, H., Wang, Y., Li, X., Zhu, J.K., et al. (2005). Disruption of the cellulose synthase gene, AtCesA8/IRX1, enhances drought and osmotic stress tolerance in Arabidopsis. Plant J 43(2), 273–283. doi: 10.1111/j.1365-313X.2005.02452.x.

de Longevialle, A.F., Small, I.D., and Lurin, C. (2010). Nuclearly encoded splicing factors implicated in RNA splicing in higher plant organelles. Mol Plant 3(4), 691–705. doi: 10.1093/mp/ssq025.

Edgar, R., Domrachev, M., and Lash, A.E. (2002). Gene Expression Omnibus: NCBI gene expression and hybridization array data repository. Nucleic Acids Res 30(1), 207–210. doi: 10.1093/nar/30.1.207.

Guo, W., Tzioutziou, N.A., Stephen, G., Milne, I., Calixto, C.P., Waugh, R., et al. (2021). 3D RNA-seq: a powerful and flexible tool for rapid and accurate differential expression and alternative splicing analysis of RNA-seq data for biologists. RNA Biol 18(11), 1574–1587. doi: 10.1080/15476286.2020.1858253.

Hernandez-Blanco, C., Feng, D.X., Hu, J., Sanchez-Vallet, A., Deslandes, L., Llorente, F., et al. (2007). Impairment of cellulose synthases required for Arabidopsis secondary cell wall formation enhances disease resistance. Plant Cell 19(3), 890–903. doi: 10.1105/tpc.106.048058.

Hu, C., Elias, E., Nawrocki, W.J., and Croce, R. (2023). Drought affects both photosystems in Arabidopsis thaliana. New Phytol 240(2), 663–675. doi: 10.1111/nph.19171.

Huang da, W., Sherman, B.T., and Lempicki, R.A. (2009). Bioinformatics enrichment tools: paths toward the comprehensive functional analysis of large gene lists. Nucleic Acids Res 37(1), 1–13. doi: 10.1093/nar/gkn923.

Ishizaki, Y., Tsunoyama, Y., Hatano, K., Ando, K., Kato, K., Shinmyo, A., et al. (2005). A nuclear-encoded sigma factor, Arabidopsis SIG6, recognizes sigma-70 type chloroplast promoters and regulates early chloroplast development in cotyledons. Plant J 42(2), 133–144. doi: 10.1111/j.1365-313X.2005.02362.x.

Kim, J.M., To, T.K., Matsui, A., Tanoi, K., Kobayashi, N.I., Matsuda, F., et al. (2017). Acetate-mediated novel survival strategy against drought in plants. Nat Plants 3, 17097. doi: 10.1038/nplants.2017.97.

Kumar, M., Atanassov, I., and Turner, S. (2017). Functional Analysis of Cellulose Synthase (CESA) Protein Class Specificity. Plant Physiol 173(2), 970–983. doi: 10.1104/pp.16.01642.

Lawson, T., and Blatt, M.R. (2014). Stomatal size, speed, and responsiveness impact on photosynthesis and water use efficiency. Plant Physiol 164(4), 1556–1570. doi: 10.1104/pp.114.237107.

Li, S., He, X., Gao, Y., Zhou, C., Chiang, V.L., and Li, W. (2021). Histone Acetylation Changes in Plant Response to Drought Stress. Genes (Basel) 12(9). doi: 10.3390/genes12091409.

Luo, M., Cheng, K., Xu, Y., Yang, S., and Wu, K. (2017). Plant Responses to Abiotic Stress Regulated by Histone Deacetylases. Front Plant Sci 8, 2147. doi: 10.3389/fpls.2017.02147.

Malbert, B., Rigaill, G., Brunaud, V., Lurin, C., and Delannoy, E. (2018). Bioinformatic Analysis of Chloroplast Gene Expression and RNA Posttranscriptional Maturations Using RNA Sequencing. Methods Mol Biol 1829, 279–294. doi: 10.1007/978-1-4939-8654-5_19.

Mao, J., Zhang, Y.C., Sang, Y., Li, Q.H., and Yang, H.Q. (2005). From The Cover: A role for Arabidopsis cryptochromes and COP1 in the regulation of stomatal opening. Proc Natl Acad Sci U S A 102(34), 12270–12275. doi: 10.1073/pnas.0501011102.

Medrano, H., Escalona, J.M., Bota, J., Gulias, J., and Flexas, J. (2002). Regulation of photosynthesis of C3 plants in response to progressive drought: stomatal conductance as a reference parameter. Ann Bot 89 Spec No(7), 895–905. doi: 10.1093/aob/mcf079.

Moazzam-Jazi, M., Ghasemi, S., Seyedi, S.M., and Niknam, V. (2018). COP1 plays a prominent role in drought stress tolerance in Arabidopsis and Pea. Plant Physiol Biochem 130, 678–691. doi: 10.1016/j.plaphy.2018.08.015.

Nunez-Vazquez, R., Desvoyes, B., and Gutierrez, C. (2022). Histone variants and modifications during abiotic stress response. Front Plant Sci 13, 984702. doi: 10.3389/fpls.2022.984702.

Patro, R., Duggal, G., Love, M.I., Irizarry, R.A., and Kingsford, C. (2017). Salmon provides fast and bias-aware quantification of transcript expression. Nat Methods 14(4), 417–419. doi: 10.1038/nmeth.4197.

Perrella, G., Lopez-Vernaza, M.A., Carr, C., Sani, E., Gossele, V., Verduyn, C., et al. (2013). Histone deacetylase complex1 expression level titrates plant growth and abscisic acid sensitivity in Arabidopsis. Plant Cell 25(9), 3491–3505. doi: 10.1105/tpc.113.114835.

Pesaresi, P., Masiero, S., Eubel, H., Braun, H.P., Bhushan, S., Glaser, E., et al. (2006). Nuclear photosynthetic gene expression is synergistically modulated by rates of protein synthesis in chloroplasts and mitochondria. Plant Cell 18(4), 970–991. doi: 10.1105/tpc.105.039073.

Seleiman, M.F., Al-Suhaibani, N., Ali, N., Akmal, M., Alotaibi, M., Refay, Y., et al. (2021). Drought Stress Impacts on Plants and Different Approaches to Alleviate Its Adverse Effects. Plants (Basel) 10(2). doi: 10.3390/plants10020259.

Singh, B., Kukreja, S., and Goutam, U. (2018). Milestones achieved in response to drought stress through reverse genetic approaches. F1000Res 7, 1311. doi: 10.12688/f1000research.15606.1.

Taylor, N.G., Howells, R.M., Huttly, A.K., Vickers, K., and Turner, S.R. (2003). Interactions among three distinct CesA proteins essential for cellulose synthesis. Proc Natl Acad Sci U S A 100(3), 1450–1455. doi: 10.1073/pnas.0337628100.

Ueda, M., Matsui, A., Nakamura, T., Abe, T., Sunaoshi, Y., Shimada, H., et al. (2018). Versatility of HDA19-deficiency in increasing the tolerance of Arabidopsis to different environmental stresses. Plant Signal Behav 13(8), e1475808. doi: 10.1080/15592324.2018.1475808.

Wilson, P.B., Estavillo, G.M., Field, K.J., Pornsiriwong, W., Carroll, A.J., Howell, K.A., et al. (2009). The nucleotidase/phosphatase SAL1 is a negative regulator of drought tolerance in Arabidopsis. Plant J 58(2), 299–317. doi: 10.1111/j.1365-313X.2008.03780.x.

Xu, D., Dhiman, R., Garibay, A., Mock, H.P., Leister, D., and Kleine, T. (2020a). Cellulose defects in the Arabidopsis secondary cell wall promote early chloroplast development. Plant J 101(1), 156–170. doi: 10.1111/tpj.14527.

Xu, D., Leister, D., and Kleine, T. (2020b). VENOSA4, a Human dNTPase SAMHD1 Homolog, Contributes to Chloroplast Development and Abiotic Stress Tolerance. Plant Physiol 182(2), 721–729. doi: 10.1104/pp.19.01108.

Xu, D., Marino, G., Klingl, A., Enderle, B., Monte, E., Kurth, J., et al. (2019). Extrachloroplastic PP7L Functions in Chloroplast Development and Abiotic Stress Tolerance. Plant Physiol 180(1), 323–341. doi: 10.1104/pp.19.00070.

Xu, D., Tang, Q., Xu, P., Schaffner, A.R., Leister, D., and Kleine, T. (2023). Response of the organellar and nuclear (post)transcriptomes of Arabidopsis to drought. Front Plant Sci 14, 1220928. doi: 10.3389/fpls.2023.1220928.

Zhang, R., Calixto, C.P.G., Marquez, Y., Venhuizen, P., Tzioutziou, N.A., Guo, W., et al. (2017). A high quality Arabidopsis transcriptome for accurate transcript-level analysis of alternative splicing. Nucleic Acids Res 45(9), 5061–5073. doi: 10.1093/nar/gkx267.

