## Supplementary for "Identification of a highly drought-resistant *pp7l hda6* mutant"

### 1 Supplementary Figures and Tables

#### 1.1 Supplementary Figures

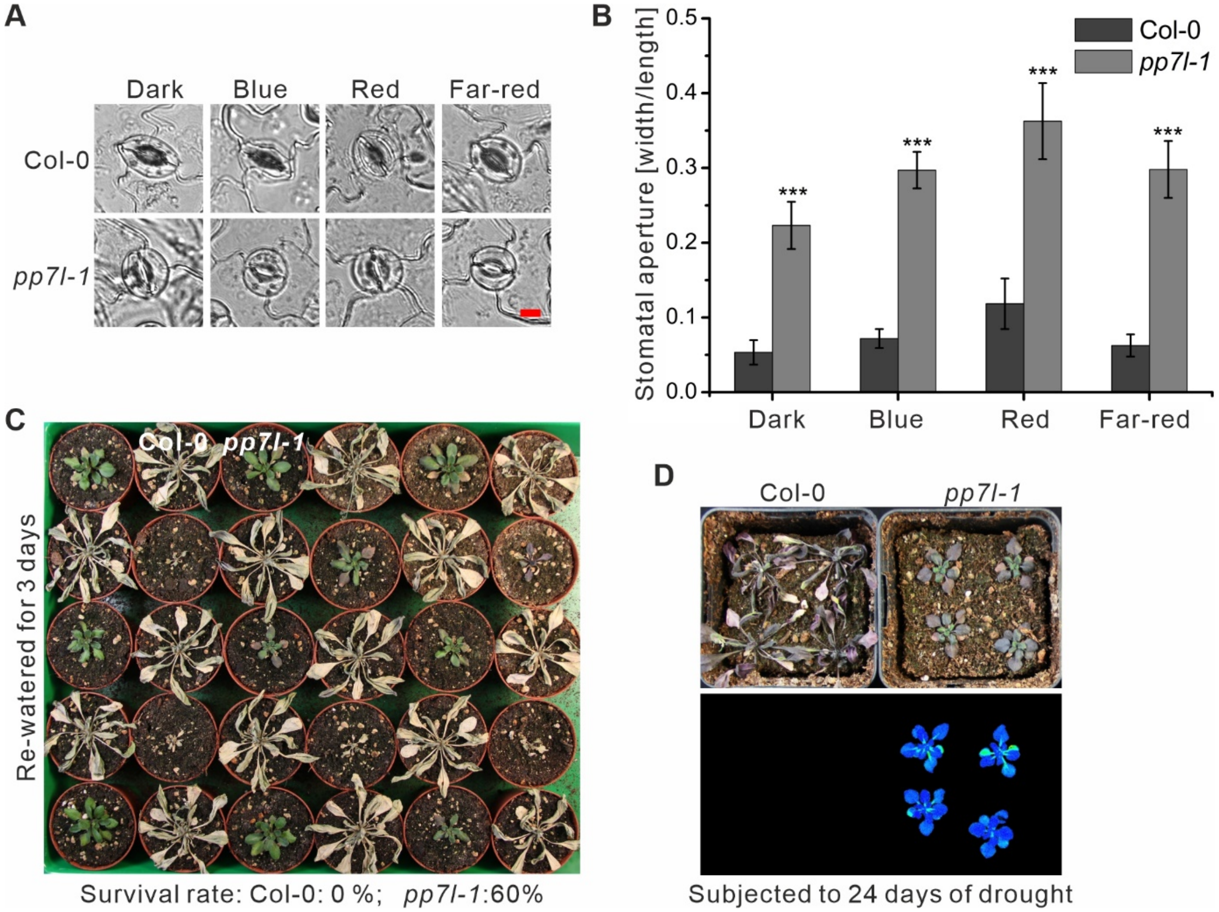


**Supplementary Figure 1.** The *pp7l* mutant can survive long periods of drought stress and behavior of stomata of Col-0 and *pp7l-1* plants in darkness, under different lighting conditions and under drought. **(A)** The micrographs show stomatal opening in leaves of Col-0 and *pp7l* mutant plants grown in the dark, or subjected to 30 μmol $m^{-2}$ $s^{-1}$ blue light, 50 μmol $m^{-2}$ $s^{-1}$ red light or 50 μmol $m^{-2}$ $s^{-1}$ far-red light. Bar =10 μm. **(B)** Analysis of stomatal aperture morphology in leaves of Col-0 and *pp7l-1* grown as in in (A). Student's t test, ****P* < 0.001. **(C)** Three-week-old plants were subjected to drought stress by withholding water for 18 days and were then re-watered for 3 days. **(D)** Phenotypes and Imaging PAM pictures of three-week-old plants that were subjected to drought stress by withholding water for 24 days.

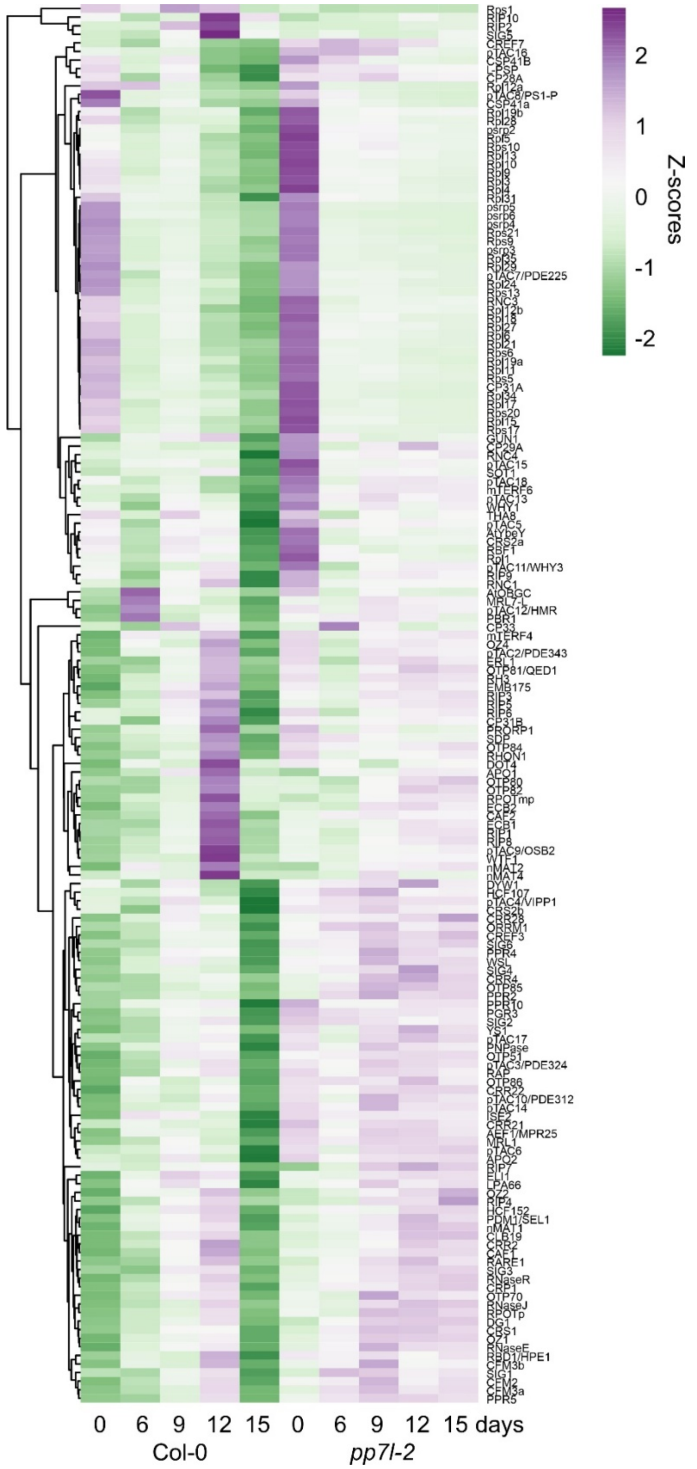


**Supplementary Figure 2.** Hierarchical clustering of transcript accumulation of nuclear genes encoding proteins involved in chloroplast gene expression. Plotting their Z-scores and sorting the transcripts into clusters according to their expression performance under drought, resulted in two clusters.

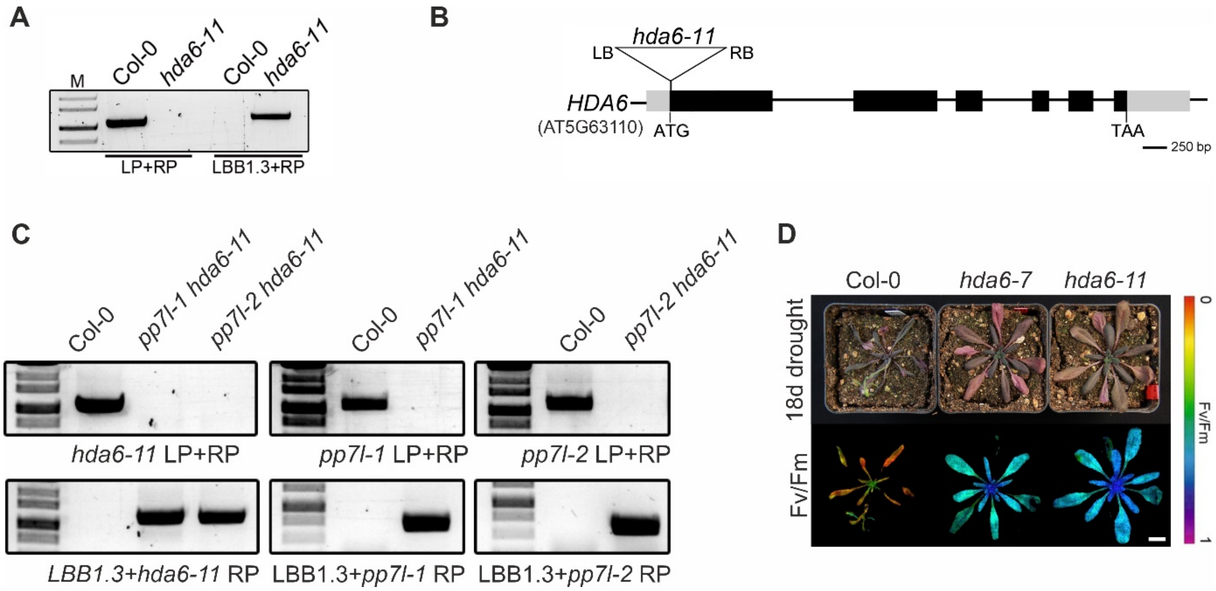


**Supplementary Figure 3.** Identification of the *pp7l-1 hda6-11* T-DNA insertion mutant. **(A)** Confirmation and identification of homozygous T-DNA insertion in the *hda6-11* mutant line. The gene-specific left and right primers (LP and RP) were used for amplification of sequences around the T-DNA insertion, and RP was used together with the T-DNA left border primer (LBB1.3) for verification of the T-DNA insertion. **(B**) Schematic representation and T-DNA tagging of the *HDA6* (*AT5G63110*) locus. Exons (black boxes), introns (black lines) and the 5' and 3' UTRs (grey boxes) are shown. Numbers are given relative to the start codon ATG. Locations and orientation of T-DNA insertions are indicated, as deduced from RP + LB PCR products shown in (A)**,** which were subsequently sequenced. Note that the insertions are not drawn to scale. **(C)** Genotyping of *pp7l-1 hda6-11* and *pp7l-2 hda6-11* mutants. PCR was performed with genomic DNA prepared from the respective plants, and PCR was performed with primer sets indicated below the gel picture. **(D)** Drought tolerance of two different *hda6* mutants compared with Col-0. Photographs were taken after 18 days of drought exposure. Bar = 1 cm.

### 1.2 Supplementary Tables

**In separate files:**

**Supplementary Table 1.** Sequencing depth of RNA-Seq experiments. RNA sequencing (RNA-Seq) was performed with RNA isolated from 3-week-old *pp7l* plants grown under optimal conditions (time-point 0 days; 0d) and after water had been withheld for 6, 9, 12 and 15 days, respectively (6d, 9d, 12d and 15d).

**Supplementary Table 2.** Abundance of chloroplast transcripts. $Log_2$ fold changes of transcripts encoded by the chloroplast genome. Note that analysis of tRNAs was excluded.

**Supplementary Table 3.** Genes differentially expressed under drought stress. RNA sequencing (RNA-Seq) was performed with RNA isolated from 3-week-old Col-0 (Col) (Xu et al., 2023) and *pp7l* (this publication) plants grown under optimal conditions (time-point 0 days; 0d) and after water had been withheld for 6, 9, 12 and 15 days, respectively (6d, 9d, 12d and 15d).
